# Systematic Reconstruction of Autism Biology with Multi-Level Whole Exome Analysis

**DOI:** 10.1101/052878

**Authors:** Weijun Luo, Chaolin Zhang, Cory R. Brouwer

## Abstract

Whole exome/genome studies on autism spectrum disorder (ASD) identified thousands of variants, yet not a coherent and systematic disease mechanism. We conduct novel integrated analyses across multiple levels on ASD exomes. These mutations do not recur or replicate at variant level, but significantly and increasingly so at gene and pathway level. Genetic association reveals a novel gene+pathway dual-hit model, better explaining ASD risk than the well-accepted mutation burden model.

In multiple analyses with independent datasets, hundreds of variants or genes consistently converge to several canonical pathways. Unlike the reported gene groups or networks, these pathways define novel, relevant, recurrent and systematic ASD biology. At sub-pathway level, most variants disrupt the pathway-related gene functions, and multiple interacting variants spotlight key modules, e.g. cAMP second-messenger system and mGluR signaling regulation by GRK in synapses. At super-pathway level, these distinct pathways are highly interconnected, and further converge to a few biology themes, i.e. synaptic function, morphology and plasticity. Therefore, ASD is a not just multi-genic but a multi-pathway disease.

## Introduction

Autism spectrum disorder (ASD) covers a range of complex genetic diseases. Genome wide molecular profiling is a proven strategy for complex disease studies. Indeed, thousands of genomic variants or loci have been identified as potential ASD causes^1-12^. These results confirm the genetic complexity of ASD and provide valuable biological insights. Yet greater challenge remains: how to turn these enormous datasets into solid and systematic understanding of the disease mechanism, i.e. biologically relevant molecular pathways, not just a list of associated genes, their groups or networks? In fact, this is the common problem remaining for all genome wide studies or complex diseases, not just ASD.

Recently, two whole exome studies under two consortia, the Simons Simplex Collection (SSC)^10,13^ and the Autism Sequencing Consortium (ASC)^11,14^, analyzed thousands of ASD families or cases-controls producing a vast amount of genetic data. These efforts identified thousands of rare mutations and firmly established their roles in ASD^15^. Because these variants rarely recur, major challenges remain as to: 1) evaluate the disease association of individual variants; 2) pinpoint most driver events from a huge pool of passengers; 3) replicate independent studies, or 4) verify their results systematically. Despite the important and inspiring discoveries, a coherent and systematic understanding of autism biology has not been achieved with these enormous studies^16^.

To address these challenges, we devised a novel integrated analysis across multiple levels, i.e. variant, gene and pathway levels. This multi-level approach has major advantages over the classical one-level approach. First, it produces more informative, systematic, holistic genetic understanding. Second, multiple-level/angle screenings of the same data is more rigorous, reaches more robust conclusions. Third, it provides more sophisticated and relevant classification and prioritization of *de novo* (DN) mutations and redefines recurrence across different levels, which makes novel and powerful analyses possible with rare events.

We applied this approach to both SSC^10^ and ASC^11^ whole exome studies, and identified hundreds of potential causal mutations. We quantified and identified substantial consistence both within and between studies, revealing a sequential convergence from variant, gene to pathway level. We deeply dissected ASD genetic association, and built a novel and more inclusive gene+pathway dual-hit model which could be generalizable to CNV or GWAS data. We reconstructed novel, replicable, systematic and multiscale molecular mechanisms for ASD. They provide solid and actionable molecular roadmaps for the development of effective and personalized ASD diagnostics and therapeutics. Note this multi-level integrated analysis is generally useful for other complex diseases, problems and genomic studies.

## Sequential convergence from variant, gene to pathway level

For multi-level analysis, we first selected ASD related DN mutations, genes and pathways in probands or cases in SSC or ASC studies as described in Methods. The results are listed in Table 1 and Table S4. We applied the same selection procedure to siblings in SSC for control.

**Table 1.** Significant pathways selected from SSC and ASC exome data. Columns *j* and *k* are the marginal and conditional counts of selected genes. These pathways are likely drivers or disease causing pathways due to the special analysis procedure (Methods). See Table S4 for full lists of selected variants and genes.

| SSC Pathways | p.val | q.val | p.con | q.con | *j* | *k* | Reference |
| --- | --- | --- | --- | --- | --- | --- | --- |
| hsa00310 Lysine degradation | $8.0 \times 10^{-05}$ | 0.02 | $8.0 \times 10^{-05}$ | 0.02 | 8 | 8 | <sup>1,2</sup> |
| hsa04727 GABAergic synapse | $4.9 \times 10^{-03}$ | 0.31 | $3.9 \times 10^{-03}$ | 0.03 | 8 | 8 | <sup>3,4</sup> |
| hsa04974 Protein digestion and absorption | $1.5 \times 10^{-02}$ | 0.35 | $9.7 \times 10^{-03}$ | 0.07 | 7 | 7 | <sup>5</sup> |
| hsa04310 Wnt signaling pathway | $7.9 \times 10^{-03}$ | 0.31 | $8.5 \times 10^{-03}$ | 0.08 | 10 | 9 | <sup>6,7</sup> |
| hsa04810 Regulation of actin cytoskeleton | $2.5 \times 10^{-02}$ | 0.35 | $6.3 \times 10^{-03}$ | 0.06 | 12 | 12 | <sup>8,9</sup> |
| hsa04010 MAPK signaling pathway | $2.1 \times 10^{-02}$ | 0.35 | $9.1 \times 10^{-03}$ | 0.07 | 14 | 10 | <sup>10,11</sup> |
| hsa04530 Tight junction | $1.7 \times 10^{-02}$ | 0.35 | $6.6 \times 10^{-02}$ | 0.17 | 9 | 5 | <sup>12,13</sup> |
| hsa04713 Circadian entrainment | $7.7 \times 10^{-03}$ | 0.31 | $6.6 \times 10^{-02}$ | 0.14 | 8 | 3 | <sup>14,15</sup> |
| hsa04724 Glutamatergic synapse | $2.1 \times 10^{-02}$ | 0.35 | $1.9 \times 10^{-01}$ | 0.19 | 8 | 2 | <sup>16,17</sup> |

**Table 1. Significant pathways selected from SSC and ASC exome data.**
| ASC Pathways | p.val | q.val | p.con | q.con | *j* | *k* | Reference |
| --- | --- | --- | --- | --- | --- | --- | --- |
| hsa00310 Lysine degradation | $6.2 \times 10^{-05}$ | 0.01 | $6.2 \times 10^{-05}$ | 0.01 | 5 | 5 | <sup>1,2</sup> |
| hsa04713 Circadian entrainment | $1.5 \times 10^{-03}$ | 0.15 | $9.9 \times 10^{-04}$ | 0.01 | 5 | 5 | <sup>14,15</sup> |
| hsa04976 Bile secretion | $3.5 \times 10^{-03}$ | 0.18 | $8.4 \times 10^{-03}$ | 0.06 | 4 | 3 | <sup>5</sup> |
| hsa04727 GABAergic synapse | $7.8 \times 10^{-03}$ | 0.21 | $4.4 \times 10^{-02}$ | 0.12 | 4 | 2 | <sup>3,4</sup> |
| hsa04010 MAPK signaling pathway | $7.1 \times 10^{-03}$ | 0.21 | $6.7 \times 10^{-02}$ | 0.07 | 7 | 4 | <sup>10,11</sup> |

ASD mutations from different cohorts do not replicate at variant level, but increasingly do so at the gene and pathway level (Fig. 1a). At variant level, 0 vs 1 of the 3348 ASC-selected variants replicate in the 1213 SSC-selected vs 3392 SSC-considered lists (p-value=0.74). At gene level, 42 vs 60 of the 182 ASC-selected genes are replicated in the 540 SSC-selected vs 1083 SSC-considered lists (p-value=9.3×10^−4^). At pathway level, 4 vs 5 of the 5 ASC-selected pathways are replicated in the 9 SSC-selected vs 199 SSC-considered lists (p-value=9.7×10^−6^). Although the background (considered) space collapses across the 3 levels as expected, the overlap ratio between studies keeps increasing. Therefore, ASD mutations show multi-level sequential convergence. In opposite, mutations from probands and siblings in the same SSC cohort do not or rarely replicate at all 3 levels, and do not converge (Fig. 1b).

**Figure 1.**
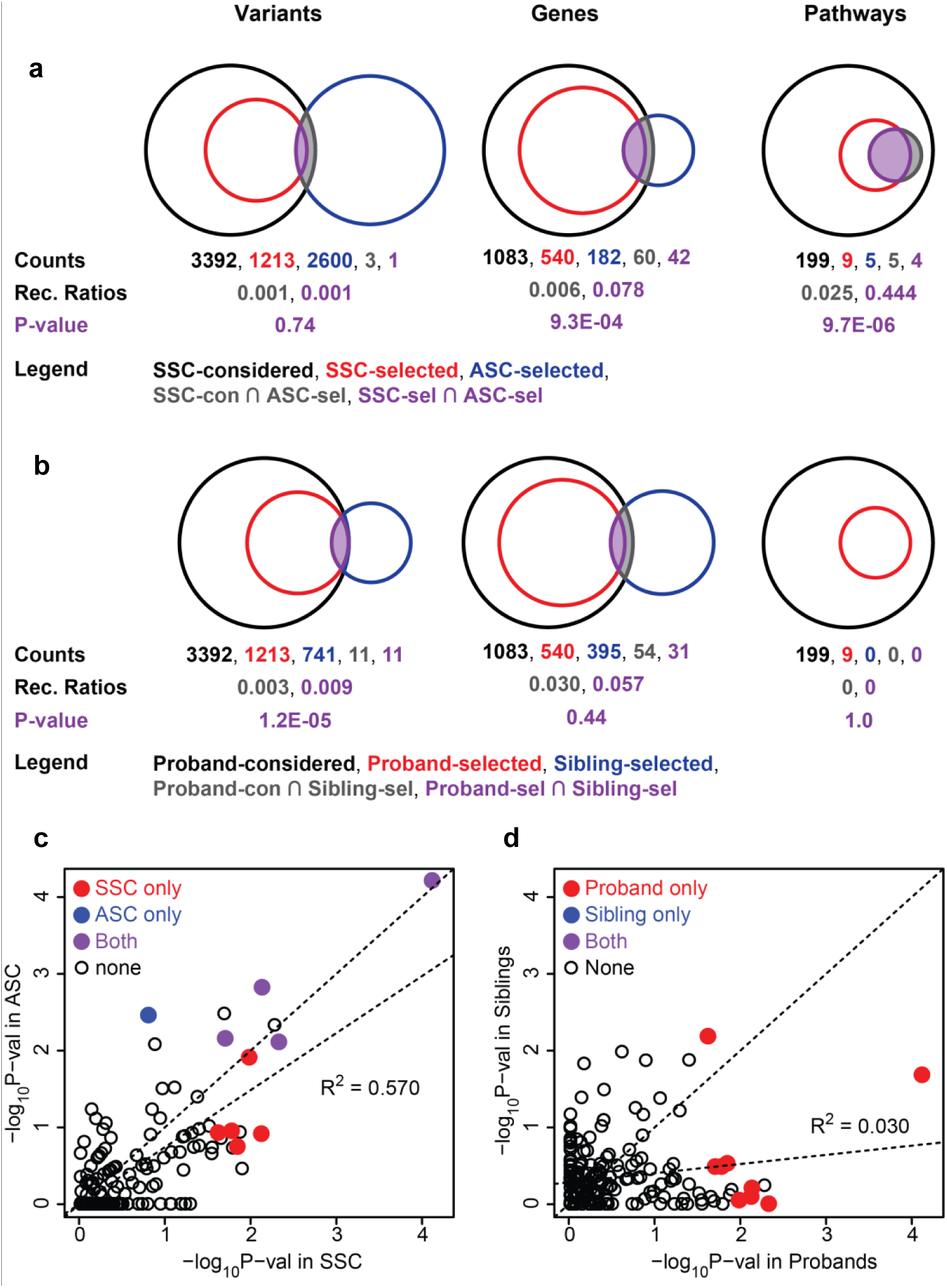
Multi-level comparison of DN mutations between or within ASD whole exome studies. Venn diagrams and test statistics on overlap a) between SSC and ASC ASD cohorts, and b) between probands and siblings of SSC, at different levels. Correlation of pathway analysis statistics c) between SSC and ASC ASD cohorts, and d) between probands and siblings of SSC. Term “considered” or “selected” refers to items before or after selection process at each level (Methods). See Table S4 for full lists of selected variants and genes used in the analysis.

Besides direct replication, pathway level analysis show extra reproducability (Fig. 1c-d). Pathway analysis statistics (-log_10_ P-val) are highly correlated between SSC and ASC (R^2^=0.570), but not so (R^2^=0.030) between probands and siblings in SSC. The actual replicability between studies should be even higher given that 1) the genetic background is much more divergent between different cohorts than between probands and siblings in the same cohort; 2) the exome-seq assay and raw data processing procedures differ between the two studies.

ASD mutations within the same cohort do not recur at variant level, but highly and increasingly so at gene and pathway level (Fig. S1a). The analysis was done with available data from SSC^10^. At variant level, 0 of 1213 selected vs 5 of the 3392 considered variants are recurrent (p-value=1.0). At gene level, 107 of 540 selected vs 110 of the 1083 considered genes are recurrent (p-value=5.0×10^−31^). At pathway level, 9 of 9 selected vs 22 of the 199 considered pathways are recurrent (p-value=8.9×10^−9^). In opposite, mutations in siblings in the same SSC cohort do not recur or less so at all 3 levels (Fig. S1b).

Gene and pathway level analysis show extra evidence of recurrence. As described above, no variants are recurrent literally, but 240 variants come from recurrent genes in probands. These gene-level recurrent variants are enriched in both the selected genes and pathways (Fig. S1c). In selected genes, such recurrent events are 31.9 and 2.53 times enriched vs in other genes and in siblings (p-vals <0.001). In selected pathways, recurrent events are 2.08 and 3.21 times enriched vs outside the pathways and in siblings (p-vals <0.001). For siblings, the selected genes but not the selected pathways are enriched for recurrent variants.

The higher-level recurrence and replication between SSC and ASC probands but not for SSC siblings: 1) indicates that our multi-level approach is both sensitive and selective; 2) suggests our results are likely true and general.

## Autism genetic association dissected across multiple levels

In this and following sections, we work with the SSC data only unless noted otherwise. The study is well controlled with simple data structure^10^, ideal for association and function analysis. For association analysis, we use all DN variants for testing power. For pathway and function analysis, we focus on validated variants only (Methods). With no recurrence or annotation, the DN events tell little on ASD genetics at variant level alone. To fully dissect the ASD genetic association, we take their gene level and pathway level effects into account. Indeed, variant effects at these two levels largely determine its association with ASD: 1) whether (and how much) the variant disrupts genes; 2) whether (and potentially how much) it hits the selected pathways.

Probands have more variants in general, particularly those disrupt genes and hit selected pathways. Probands have 55% more LGD or Likely Gene Disrupting (0.175 vs 0.113, p=2.1×10^−6^) and 11% more missense variants (0.667 vs 0.601, p=2.2×10^−2^) than siblings, but only 6% more silent variants (0.515 vs 0.484, p=7.8×10^−2^) (Fig. S2 row 1). This is consistent with the original analysis^10^. In addition, they have 39% more variants within selected pathways, but only 12% more outside (Fig. 2 row 1). In other words, most of the differences between probands and siblings fall in LGD and missense categories within selected pathways. This difference is well exhibited by variant distributions in Wnt and synapse pathways (Fig. S5-7).

**Figure 2.**
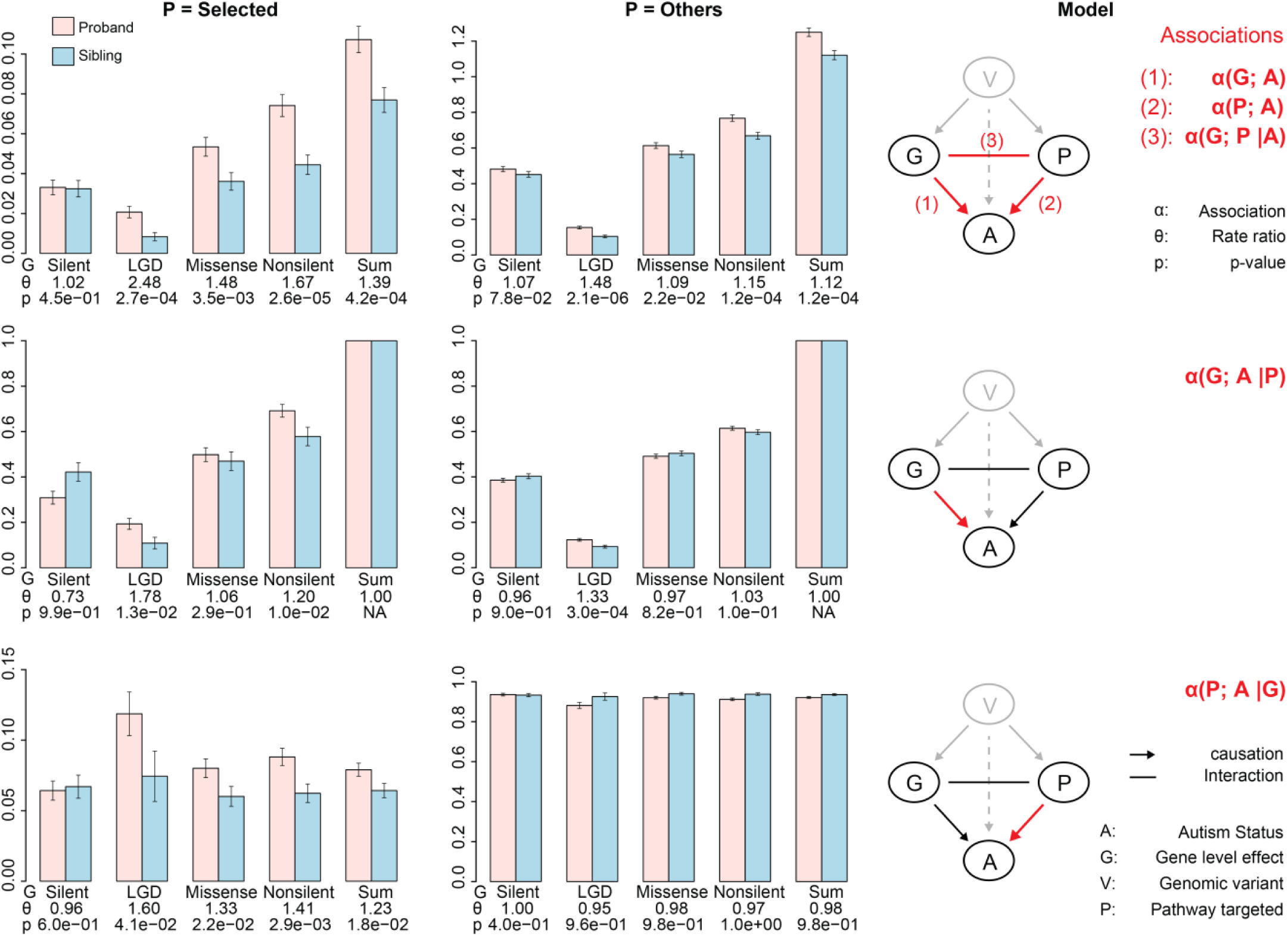
Autism genetic association analysis across variant, gene and pathway levels with the SSC exome mutation data. Column 1 and 2: DN event rates and association test by gene and pathway level effects; column 3: A descriptive model for autism genetic association. Rows are marginal (Row 1) and conditional (Row 2-3) association tests and statistics, and corresponding model representations. Variants are grouped based on gene-level effects (G): Silent, Missense, LGD and Nonsilent (Missense + LGD) (details in Methods), and pathway level effects (P): hitting Selected pathways or Others. The associations marked in column 3 should be taken as the theme for corresponding row. Notation α(G; A) and α(G; A |P) are read as marginal association between G and A, and their conditional association given P. The association can be measured by rate difference (over noise), rate ratio (α) or log α. P-values comes from the rate difference tests. Error bars represent standard error of the mean (SEM).

Proband variants are more (likely, frequently) gene disrupting. As described above, probands have more events in LGD and missense categories and in general. After adjusting for the event numbers per pathway assignment or in total (row 2 in Fig. 2 and Fig. S2), probands still have 78%, 33% and 37% higher LGD within selected pathways, outside and together (p= 1.3×10^−2^, 3.0×10^−4^ and 2.8×10^−5^respectively). This difference is well exhibited by in Wnt and synapse pathways (Fig. S5-7). The missense variant ratios are similar in probands and siblings (row 2 in Fig. 2 and Fig. S2). However, greater portions of missense variants are gene damaging in probands vs siblings (0.58 vs 0.43 and 0.50 vs 0.47 within and outside selected pathways, p<0.05 and 0.1 respectively) as predicted by SIFT^17^ (Fig. 4a). In addition, more missense variants in selected pathways hit a functional domain in probands vs siblings (0.747 vs 0.558, p-val<0.05) (Fig. 4b), especially in Wnt and synapse pathways (Fig. S8-9, and more details in Supplementary Text Section 4).

Proband variants are more (likely, frequently) pathway hitting. Probands have higher absolute event rates than siblings in selected pathways (0.11 vs 0.08, p= 4.2×10^−4^), especially in LGD (0.021 vs 0.008, p= 2.7×10^−4^) and missense (0.053 vs 0.036, p= 3.5×10^−3^) categories, but not the silent category (0.033 vs 0.032, p=0.45). After adjusted for event numbers within each category or in total, probands still have consistently higher pathway event rates than siblings for both LGD (0.12 vs 0.07, p= 4.1×10^−2^) and missense categories (0.08 vs 0.06, p= 2.2×10^−2^), but not for the silent category (0.06 vs 0.07, p= 0.60) (row 3 in Fig. 2).

We proposed a gene+pathway dual-hit (or two-factor) model for ASD genetic association based on our results above (Fig. 2 column 3): disrupting effect on target genes (G) and hitting the relevant pathways or not (P). These two factors have significant association with ASD both marginally and conditionally as described above. In our model, variant load/burden per person (V) becomes less relevant and marked as hidden. Because the extra variants mostly fall into the gene disrupting and pathway hitting categories (Fig. 2 row 1, described above).

There is also significant interaction between gene and pathway factors (Fig. 2 row 1). Probands and siblings have the biggest differences in variants that are both gene disrupting and pathway hitting. The differences diminish outside the pathways or disappear completely in the silent category. Indeed, this interaction is significant, as indicated by significant overrepresentation in LGD hitting the pathways in probands (52 occurred vs 34.6 expected events, p=0.001, Table S1).

What we proposed is essentially a Noisy-AND model, i.e. risk genetic variant tend to be both gene disrupting AND pathway hitting. The model is Noisy because our knowledge on gene disrupting and pathway assignment is incomplete or penetrance incomplete.

We also estimate the prevalence of DN events with different gene and pathway level effects (Fig. S3). These statistics are similar to per patient variant burden stats (Fig. 2 row 1) and consistent with our two-factor genetic model for ASD (Fig. 2 row 1). With the DN variants alone, this model explains at least 5% (2.8% within and 2.2% outside selected pathways) of all ASD cases (Fig. S3). Although consistent with the LoF mutation contribution in the ASC study^11^, this is likely a substantial under-estimation, since not all variants are called and not all genes in the relevant pathways are known. In addition, when other types of variants (CNV, common variants or transmitted/inherited variants) are considered, this model can be generic and more descriptive (more in Supplementary Text Section 2).

## Pathways of DN events, integrated molecular mechanism

Selected by our special pathway-level testing procedure, these pathways form a novel, coherent yet non-redundant set of ASD disease mechanisms (Table 1). <u>These pathways are novel in multiple aspects: 1) first time report the target pathway is involved in ASD (Actin, MAPK, T-junction), 2) there are some evidence in literature, but this is the first report based on whole exome/genome analysis with statistical significance (Lysine, GABA, Wnt, Circ, Glut); 3) for all pathways, this is the first report with pathway graphs on detailed molecular mechanisms for ASD; 4) mostly being causal, they may also explain associated symptoms of ASD, including intellectual disability^18^ (Glut, GABA), sleeping^19^ (Circ) and digestive^20^ (digestion) problems</u>.

Importantly, the pathway graphs integrate disease variants and genes from multiple datasets: SSC^10^, ASC^11^ or SFARI Gene database^21^ (Fig. 4, Fig. S4). These pathways are likely true and primary molecular mechanism for ASD as they are consistently selected in these independent analyses. These analyses agree in details too: they frequently converge to the same genes, gene groups (nodes) or the same signaling branch in a pathway. They also complement each other. For instance, SSC and ASC data provide numerous novel ASD associated genes besides those collected in SFARI Gene.

The pathway list and data integrated pathway graphs provide abundant novel, coherent and systematic insights on ASD mechanism. We focus on three pathways for example.

### 1. Wnt signaling pathway: the canonical branch only (Fig. 4a).

All DN events from SSC and ASC, and all SFARI genes converge to the canonical Wnt pathway. However, just a few events/genes hit noncanonical Wnt pathways, which are mostly shared by the canonical branch or other selected pathways.

In addition, all aspects or steps of canonical Wnt signaling are involved in ASD (Fig. 4a). These include Wnt and co-receptor LRP5/6, messenger Dvl, key component of the destruction complex: APC, GSK3 and CKIε, other key players in β-catenin phosphorylation/ubiquitination/degradation: TBL1 in p53-induced SCF-like complex, and β-TrCP in Skp1-Cullin-F-box (SCF) E3 ubiquitin ligase complex, PS-1 (Presenilin). Finally, repressors or activators (chromatin remodelers) in β-catenin directed transcription, CHD8 (Duplin), RUVBL1 (Pontin52), CREBBP (CBP).

### 2. The whole GABAergic synapse pathway is involved in ASD

particularly the following parts (Fig. 4b):

1. GABAA receptor or signal in the postsynaptic neurons, and the negative feedback loops (Gi/o, AC) in pre-and post-synaptic neurons, and the clearance channel through GABA transporters (GATs) on the presynaptic terminal or neighboring glial cells.
2. Glial cells besides the presynaptic and postsynaptic neurons.

### 3. The whole Glutamatergic synapse pathway is involved in ASD

particularly the following parts (Fig. 4c):

1. ionotropic glutamate receptor (iGluRs, NMDARs) signal, and the postsynaptic density scaffold proteins (SHANKs etc), the consequent synapse formation and plasticity.
2. metabotropic glutamate receptors (mGluRs, mGluR1, 5, 7, 8), the coupled *G* proteins (Gs, Gi and Go) and the second messenger systems downstream (Ca2+, cAMP, DAG, IP3).
3. the inhibitory autoreceptor mechanism that suppresses excess glutamate release in presynaptic neurons (mGluR7, Gi/o, GRK, AC).
4. Glial cells besides the presynaptic and postsynaptic neurons, especially in the clearance and recycle of glutamate.

Other pathways and graphs are equally informative, many of them are also supported by literature (Table 1). For details, please check the Supplementary Text Section 3 and Fig. S4.

## Subpathway biology, coherent fine details

We analyzed the functional consequences of DN variants in selected pathways. Here we focus on missense but not LGD variants. Because the latter are highly destructive on overall protein structure and function (position insensitive), while the former are subtle and precisely tell what functions are perturbed in ASD (position sensitive).

In probands, missense variants hit the relevant functions or domains in selected pathways.

In Wnt signaling pathway, missense hit the histone acetylation domain KAT11 twice in CREBBP (CBP) gene and TIP49 domain in RUVBL1, the scaffolding domain WD40 in TBL1XR1, and the CTNNB1 binding domain in TCF7L1 (Fig. S8). In synapse pathways, the most essential players, i.e. neurotransmitter receptors, transporters and ion channels on cell membrane, are heavily targeted (Fig. 4c-d, Fig. S6-7). Missense variants hit the neurotransmitter glutamate binding domain in GRIN2B (NMDAR) gene, the 7 transmembrane region of GRM7 (mGluR7), and the ion-channel domains in GABRA1 and CACNA1C, the Sodium:neurotransmitter symporter domain in SLC6A1 (GAT) and SLC6A13 (GAT2/3), among others.

In opposite, missense variants in siblings often hit the non-functional regions or the less relevant regions or genes (Fig. S8-10).This probands-sibling difference is significant overall (Fig. 4b) and extremely so in the example pathways (Fig. S8-9 and Supplementary Text Section 4).

Autistic missense events on the same genes tend to hit residues extremely close and in the same domain. This occurs to all cases we observed in Wnt and synapse pathways (Fig. 4c-d or Fig. S8-9: ADCY5, CREBBP, SLC6A1 and SLC6A13). These data strongly suggest that missense events do not occurred in random, but precisely and consistently targeting specific risky loci for ASD (p= 0.002-0.03, Supplementary Text Section 4).

We identified subpathway clusters of missense events in probands. Each event cluster hits multiple interacting genes along the pathway. They reveal novel and critical molecular modules in ASD biology.

One cluster hit the cAMP second-messenger system^22^ in the Glutamatergic synapse pathway (Fig. 4d, Fig. S7). Two types of G proteins bind and control Adenylate cyclase (AC), Gs activates while Gi/o (Gi/Go) inhibits it (green dashed box in Fig. S7). As shown in Fig. 4d, the G-alpha domains of GNAS (Gs) and GNAO1 (Gi/o) are similar and align seamlessly in 3D^23^. They compete to bind to AC C2A domain the same way. Missense variants in these two genes both hit the G-alpha domain, which affect their binding to AC hence AC’s catalytic activity on cAMP production and downstream signal. In parallel, the two missense events on AC (ADCY5) hit its C1A domain, which perturb AC’s catalytic function too (Fig. 4d). In the direct upstream (Fig. S7), GRM5 (mGluR5) was hit by a destructive in frame deletion (K679) (Table S5). GRM7 (mGluR7) was hit by a missense at the 7 transmembrane region (Fig. 4c), which likely render it a strong antagonist of the transmembrane signal as an unbounded cytosol form.

In another cluster, GRK inhibits mGluR signaling by sequestering heterotrimeric *G* proteins. See Supplementary Text Section 4 and Fig. S11 for details.

All these subpathway level biological stories we present above reveals coherent fine details on ASD mechanism. This is consistent with yet complement to the integrated pathway graphs (Fig. 3 and Fig. S6-7).

**Figure 3.**
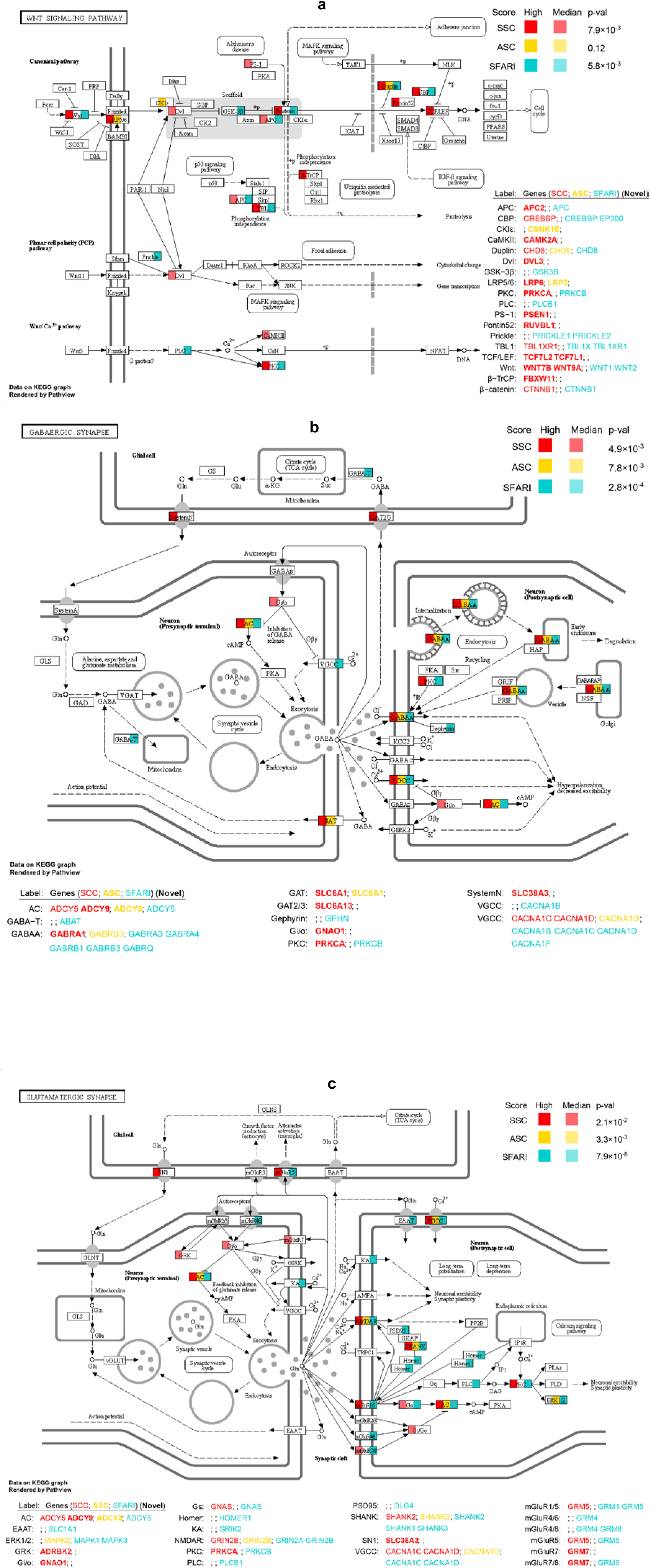
An integrated view of autism associated DN variants or genes from multiple sources in selected KEGG pathways: a) hsa04310 Wnt signaling pathway, b) hsa04727 GABAergic synapse, and c) hsa04724 Glutamatergic synapse (next page). DN variants data come from SSC and ASC studies, and reported autism genes from SFARI Gene Database. Gene level scores (Methods) are marked by color. P-values are from pathway analysis (Table 1). Data are integrated and visualized on KEGG pathway graphs using Pathview (23).

**Figure 4.**
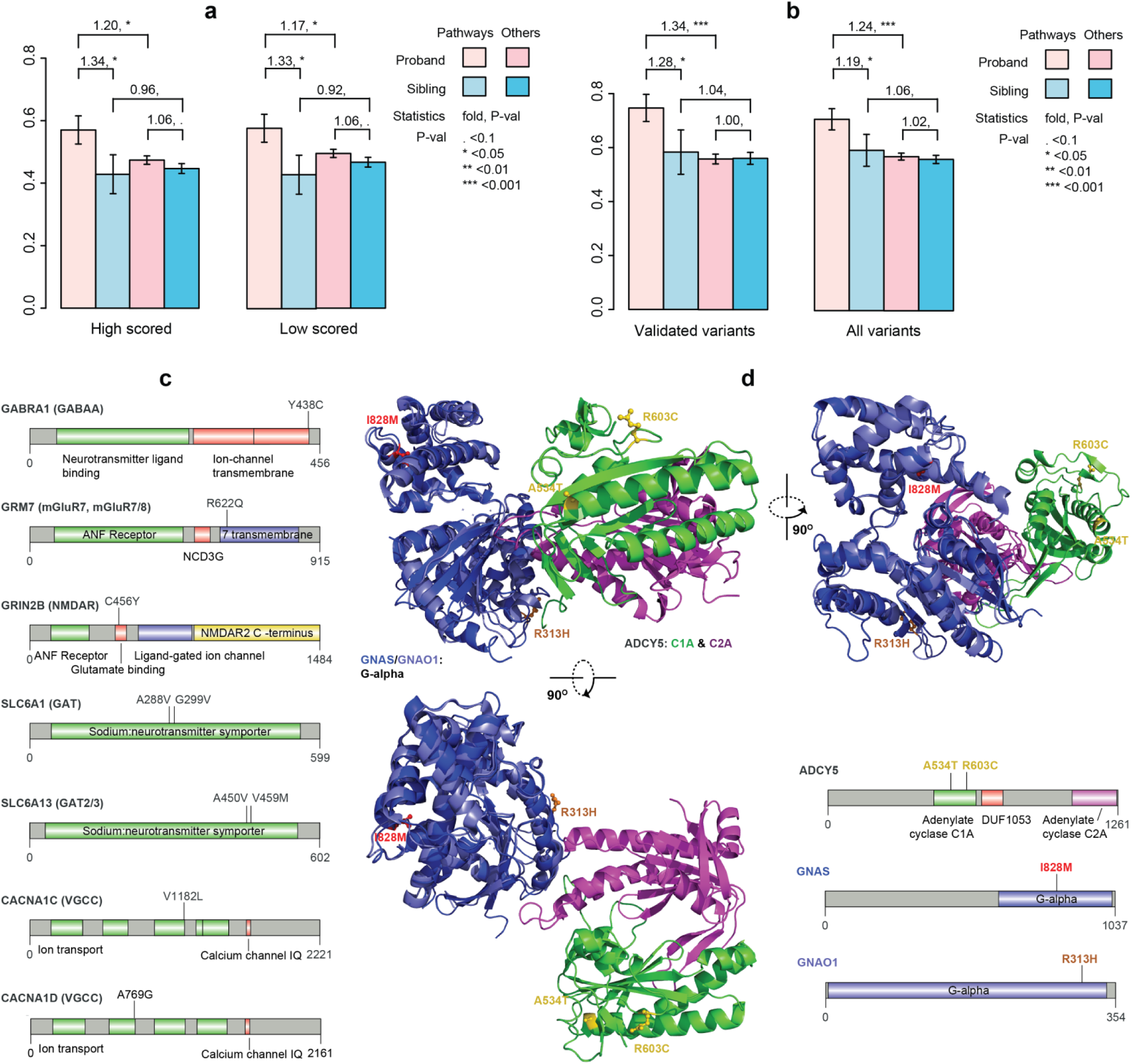
Functional consequences of autism associated missense mutations. a) ratios of damaging events as predicted by SIFT; b) ratios of events hitting a function domain defined in Pfam; c) 1D protein domain structure and missense variants of all neurotransmitter receptors, transporters and ion channel genes in synapse pathways (the bold black box nodes in pathway graphs in Figure S6-7); d) 1 and 3D protein structures and missense variants hitting the Adenylate cyclase (AC), i.e. ADCY5, and interacting G proteins, GNAS (Gs) and GNAO1 (Gi/o). The pathway context is shown in the green dashed box in in Supplementary Fig. 7. AC controls the production of cAMP second-messenger in synapse (Supplementary Text Section 4). Error bars represent standard error of the mean (SEM).

## Superpathway biology, emergent big picture

The selected pathways are distinct yet highly interconnected. For example, MAPK feed into canonical Wnt pathway and inhibit TCF/LEF dependent transcription (Fig. 3a and Fig. S4c). In addition, they also share numerous other connections. For example, Wnt and MAPK are both involved in adherens junctions and focal adhesion (Fig. S4g-4h). These commonly connected pathways are also perturbed in ASD except marginally significant (p.val=0.01-0.10, Table S2).

Two distinct biological themes or modules emerge from the selected and connected pathways (Fig. 5). Module I includes Wnt signaling, cell adhesion, junction, and cytoskeleton etc. They are involved in synapses morphology, i.e. synapse assembly and stability. Module II includes Glutamatergic synapse, GABAergic synapse, and related processes. They are involved in synapses functions, i.e. chemical and electrical signals transmission, regulations and patterns. Module 1 concerns neuronal wiring or the hardware, while module II concerns synaptic transmission or the software. These modules are distinct in topology too. Connections are dense within each module but none between them. MAPK pathway is the only bridge node and highly connected in both modules. There is 1 less prominent theme: transcription (not shown in Fig. 5). Both Wnt and MAPK pathways end at target gene transcription, which involves chromatin modification, especially histone lysine methylation branch of Lysine degradation (Table 1). We also did a parallel GO term analysis, which converges to the same set of biological themes (Supplementary text section 5 and Fig. S12).

**Figure 5.**
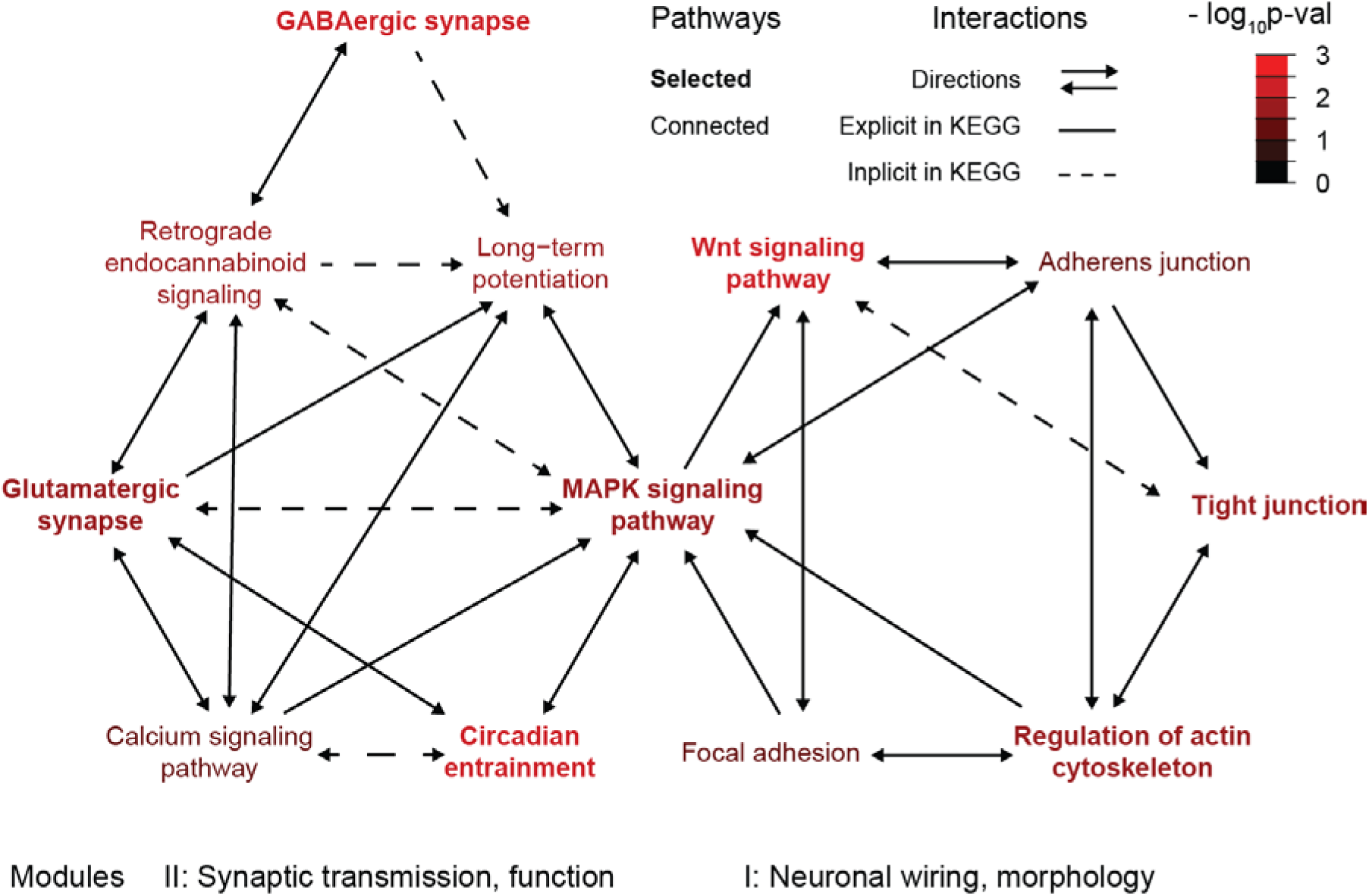
The super-pathway clusters emerged from the pathway-level analysis of SSC exome variant data. Seven selected pathways are highly interconnected and frequently connected to 5 additional pathways. All these pathways form a super-pathway level network. Two clusters or modules emerged with distinct topology and function.

All mutated pathways or functions converge to synapse biology. Either synaptic function, morphology or plasticity (as indicated by transcription^24,25^) is disrupted in these cases. Therefore, ASD is a multi-pathway disease not just multi-genic, and ultimately a synapse disease.

## Discussion

We conduct an integrated analysis on ASD exome mutations across multiple levels. These isolated and rarely occurred events are actually connected and recurrent at higher (gene and pathway) levels (Fig. 1). In the meantime, the otherwise random and divergent results become reproducible between independent studies. This cross-validation not only confirms our results but also justifies our multi-level analysis approach. This novel approach is equally applicable to other complex diseases.

We also did a multi-level association analysis, and proposed a gene+pathway dual-hit model for ASD risk (Fig. 2). The disease variants need to both: 1) disrupt the target genes; and 2) hit the relevant pathways. Variants missing either factor become no or less risky, including the silent variants in the selected pathways or variants outside the pathways. In this model, contribution of variant load/burden can be explained away hence becomes less relevant. This model likely applies to other types of genetic variants including CNV and SNP. Although this is just a descriptive model, with relevant data, it can turn into a predictive model.

We reconstruct a set of coherent and systematic molecular mechanisms for ASD (Fig. 3-5, Table 1). Importantly, we discover whole pathways or molecular systems that cause the disease, as supported by multiple independent datasets. These disease pathways not just present a catalog of ASD genetic associations (Table S5), but further connect hundreds of interacting genes and variants into a whole, dynamic multiscale system (Fig. 3-5). They reveal concrete biological mechanism, much more definitive and informative than gene networks or GO groups in literature. These results greatly advance our understanding on ASD, and provide solid guidance for the development of effective diagnosis and therapeutics on ASD.

## Methods

### Data collection and integration

The exome-seq DN variants from the SSC cohort^10^ and ASC cohort^11^ were used for this study. Please see the original publications for details of the experimental design, quality control and raw data processing. The final SSC data include 2,517 families, with 2,508 affected children, 1,911 unaffected siblings and the parents of each family. The ASC data we used consists two cohorts: one includes1,445 trios, another includes1601cases and 5397 ancestry-matched controls. The ASC paper originally included 825 trios from the SSC cohort. This overlap was intentionally excluded to create two completely independent datasets for downstream analysis and comparison.

### Variant level analysis

Variants were divided into 3 major categories based on their effects on the target genes. Silent group includes all synonymous variants and those fall in the 3'UTR, 5'UTR, intergenic, intron, and non-coding regions; Missense group include missense variants; LGD (likely gene disrupting) or LoF (lose of function) group includes exon indels (both frame-shift and no-frame-shift), nonsense, and splice-site variants. Variants are selected for gene and pathway level analyses based on a few criteria:1) LGD (or LoF) and missense only, as silent variants are usually not damaging, and have little disease association as a group (Fig. 2); 2) For SSC study^10^, we only consider validated variants, which included those experimentally verified or cross-validated or called in at least 2 of the 3 laboratories (CSHL, Yale or UW). 3) For ASC study^11^, we only consider DN variants in the trio families or those from the case-control cohorts.

### Gene level analysis

Selected variants are mapped to target genes. We select genes using the following scoring function which essentially sums up the weighted evidence for each gene.

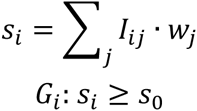

*i*: gene index, *j*: patient index, *I*, indicator on whether a selected variant occurs to the gene-patient pair, *w*, weight, *s*: score

Due to different study designs and data quality, we used slightly different criteria for the two cohorts. In SSC, we take *w*_*j*_=1/*n*_*j*_ (number of selected variants occurred to patient i) and *s*_0_=0.5, while in ASC *w*_*j*=1 and *s*_0_=2_.

### Pathway level analysis

We selected pathways enriched for the selected genes. We test for both marginal and conditional overrepresentation given the previously selected pathways. This procedure ensures that pathways selected are drivers instead of passengers, which share genes with the former.

The analysis is an application of the set theory.

*G* = {selected genes above}

*Pi* = {pathway or gene set under testing}

*Ps* = {selected pathways or gene sets}

*P*_*0*_ = {all pathways or gene sets}

*U* = | *G* ∩ *P*_*0*_ |

*V* = | *G* ∩ *P*_*0*_\*Ps* |

*X* = | *G* ∩ *Pi* |

*Y* = | *G* ∩ *Pi*\*Ps* |

For marginal significance test:

*X* = *j* ~ hyperG(*j*; |*P*_*0*_|, |*Pi*|, *U*)

*P*(*X* ≥ *j*) = Σ_*l*_ PhyperG(*j*; |*P*_*0*_|, |*Pi*|, *U*), where *l*= { *j*, *j*+1, ‥ |*P*_*0*_|}

For conditional significance test:

*X* = *k* | *Ps* ~ *Y* = *k* ~ hyperG(*k*; |*P*_*0*_\*Ps*|, |*Pi*\*Ps* |, V)

*P*(*X* ≥ *k* | *Ps*) = P(*Y* ≥ *k*) = Σ_*l*_ PhyperG(*k*; |*P*_*0*_\*Ps*|, |*Pi*\*Ps* |, V), where *l*= {*k*, *k*+1, ‥ |*P*_*0*_|}

Here hyperG is the hypergeometric distribution, and PhyperG is the standard probability mass function of the hypergeometric distribution.

The same analysis procedure was applied to KEGG pathways and GO terms. The metabolic and signaling pathways from KEGG were tested and analyzed together, and the three branches of GO, i.e. biological process (BP), cellular component (CC), molecular function (MF) were analyzed separately. We did multiple-testing correction on P-values using false discovery rate (FDR or q-value).

### Variant association

ASD DN variants can be divided into groups based on their gene-level and pathway-level effects. At gene-level, they are assigned to silent, LGD, missense or nonsilent (LGD + missense) groups, as described above. At pathway level, they either belong to the Selected pathways or Others.

The ASD association of these variant groups can be measured by rate difference (over noise), rate ratio (θ) or log θ between probands and siblings. To test the rate difference between probands and siblings, we conducted two proportion z-test for conditional rates, and two sample t-test for marginal rates. Odd ratio tests gave similar results as in our conditional rate tests, but is not suitable for marginal tests on absolute variant rates.

### Pathway data integration and visualization

Pathview package was used for pathway based data integration and visualization. Variants were first mapped to the target genes, which are then mapped and visualized onto the selected KEGG pathway graphs. In disease gene view (Fig. 3, Fig. S4), variant targeted genes from SSC and ASC, and SFARI genes are collected, integrated and shown in the relevant pathways. Different data sources were marked by colors, gene level scores by brightness, and corresponding pathway analysis p-values are also shown. In variant type views (Fig. S5-7), DN variants from SSC are project and visualized on the target pathways. Variant types or effects (LGD, missense, or silent) are marked by different colors, their corresponding event counts are also shown.

### Protein structure and function analysis

Exome variants were mapped to amino acid changes in the target protein using Bioconductor Vari ant Annotation package. 1D Linear protein domain structures were visualized using cBioPortal Mutationmapper. Protein domain data were retrieved from Pfam database, and provide updated the protein domain locations. 3D protein structure data were retrieved from the Protein Data Bank (PDB). The mapped exome variants coded into amino acid changes were then visualized with the 3D protein structure using Pymol (www.pymol.org).

